# 2018-2019 human seasonal H3N2 influenza A virus spillovers into swine with demonstrated virus transmission in pigs were not sustained in the pig population

**DOI:** 10.1101/2024.01.12.575431

**Authors:** Joshua D. Powell, Megan N. Thomas, Tavis K. Anderson, Michael A. Zeller, Phillip C. Gauger, Amy L. Vincent Baker

**Author notes:** Corresponding author: Amy Vincent Baker, Virus and Prion Research Unit, NADC, USDA-ARS, 1920, Dayton Avenue, PO Box 70, Ames, IA 50010, USA. National Animal Disease Center, 1920 Dayton Avenue, P.O. Box 70, Ames, IA 50010-0070, USA, Voice: 1-515-337-7557 • •, USDA is an Equal Opportunity Provider and Employer.

## Abstract

Human seasonal H3 3C3a clade influenza A viruses (IAV) were detected in four U.S. pigs from commercial swine farms in Michigan, Illinois, and Virginia in 2019. To evaluate the relative risk of this spillover to the pig population, whole genome sequencing and phylogenetic characterization was conducted and revealed all eight viral gene segments were closely related to 2018-2019 H3N2 human seasonal IAV. Next, a series of in vitro viral kinetics, receptor binding, and antigenic characterization studies were performed using a representative A/swine/Virginia/A02478738/2018(H3N2) (SW/VA/19) isolate. Viral replication kinetic studies of SW/VA/19 demonstrated less efficient replication curves than all ten swine H3N2 viruses tested, but higher than three human H3N2 strains. Serial passaging experiments of SW/VA/19 in swine cells did not increase virus replication, but changes at HA amino acid positions 9 and 159 occurred. In swine transmission studies, wild type SW/VA/19 was shed in nasal secretions and transmitted to all indirect contact pigs, whereas the human seasonal strain A/Switzerland/9715293/2013(H3N2) from the same 3C3a clade failed to transmit. SW/VA/19 induced minimal macroscopic and microscopic lung lesions. Collectively these findings demonstrate that these human seasonal H3N2 3C3a-like viruses did not require reassortment with endemic swine IAV gene segments, impacting virus shedding and transmission in pigs. Limited detections in the U.S. pig population in the subsequent period of time suggests a yet unknown restriction factor likely limiting the spread of these viruses in the U.S. pig population.

**IMPORTANCE:** Interspecies human-to-swine IAV transmission occurs globally and contributes to increased IAV diversity in pig populations. We present data that a swine isolate from a 2018-2019 human-to-swine transmission event was shed for multiple days in challenged and contact pigs. By characterizing this introduction through bioinformatic, molecular, and animal experimental approaches, these findings better inform animal health practices and in vaccine decision-making. Since wholly human seasonal H3N2 viruses in the U.S. were not previously identified as being transmissible in pigs (i.e. reverse zoonosis), these findings reveal the interspecies barriers for transmission to pigs may not require significant changes to all human seasonal H3N2.

## 1. INTRODUCTION

Influenza A virus (IAV) of the family Orthomyxoviridae, has a segmented negative-sense RNA genome and causes respiratory disease in numerous hosts, including humans, pigs, and birds. IAV caused four human pandemics since the early 1900’s (1); the introduction of a novel hemagglutinin (HA) from animal hosts into the human population was a hallmark of each emerging pandemic strain. While IAV pandemic events arising from animal reservoirs have been documented for more than a century, only more recently is it recognized that human seasonal IAV frequently spills over into the swine population (2, 3). As human-to-swine reverse zoonosis continually increases the diversity of swine IAV in circulation, there are now numerous genetically and antigenically diverse co-circulating IAVs. This heterogeneity has substantial economic impact due to the respiratory disease inflicted on pigs by IAV and the inability to target all circulating IAVs with a licensed multivalent vaccine. In addition to swine health, zoonotic swine IAV that spill over into humans, referred to as “variant (v)” IAV, are of public health importance, with at least 498 human cases of variant infections reported by the CDC since 2010 (https://gis.cdc.gov/grasp/fluview/Novel_Influenza.html). Consequently, identifying spillovers at the swine-human interface in both directions is critical and requires a One Health approach with collaborative support of human and animal health experts at the clinical, diagnostic and public health levels (4).

IAVs are categorized based on the hemagglutinin (HA) and neuraminidase (NA) surface glycoproteins, presently encompassing 18 HA and 11 NA subtypes (5, 6). The H1N1, H3N2 and H1N2 IAV subtypes co-circulate in the global swine population with occasional detections of H3N1. Multiple reviews summarized the genetic and antigenic relationships of the H1 and H3 swine viruses that circulate globally in pigs (7, 8). H3 viruses detected in the United States include three major swine H3 lineages that are presently sustained in the U.S. pig population. The 1990.4 H3 clade (known colloquially as cluster-IV), have been endemic since the late 1990’s (9–11). Clade 2010.1, a result of human H3N2 spillover in the early 2010’s also widely circulates (12, 13). Clade 2010.2, a lineage from a more recent human-to-swine 2016-2017 H3N2 spillover is sporadically but consistently detected (14, 15). A unifying theme within all three lineages is that shortly after human-to-swine virus introduction there was the replacement of human (H) internal virus segments with either the endemic triple-reassortant lineage that has been in swine populations since the 1990s (TRIG or T) or from the 2009 H1N1 pandemic lineage (P) through reassortment (12, 13, 16, 17). Full genome sequencing of swine isolates for the PB2, PB1, PA, NP, M and NS internal genome segments frequently detected TTTTPT and TTTPPT arrangements. Human (H) internal segments retained from human-to-swine spillovers (HHHHHH) were not previously identified in the U.S through USDA surveillance.

In 2019, four swine respiratory samples originating from Illinois, Michigan and Virginia were submitted to the Iowa State University Veterinary Diagnostic Laboratory and confirmed positive for IAV. Whole virus genome sequencing in conjunction with the USDA swine influenza surveillance program and phylogenetics performed as part of this study revealed two important facets of these viruses. First, these detections were closely related to recent seasonal human H3N2 IAV from the 3C3a clade that circulated in North America during the 2018-2019 flu season. Second, these viruses had not yet undergone reassortment with circulating swine IAV and retained their respective HHHHHH internal segments in addition to their HA and NA glycoproteins. Consequently, we set out to experimentally study a representative isolate A/Swine/Virginia/A02478738/2019(H3N2) (SW/VA/19) by characterizing the virus in swine respiratory cells *in vitro* and in a swine transmission study *in vivo*. In swine cells, SW/VA/19 viral kinetics were compared to ten different swine strains and three North American human seasonal H3N2 vaccine strains. SW/VA/19 was passaged in swine respiratory cells to evaluate survival and evolution in natural host cells. Receptor binding and antigenic testing of SW/VA/19 was also assayed and compared to recent contemporary viruses from the two major 1990.4 and 2010.1 swine H3 clades to provide further insight into the relationship currently circulating swine viruses have to these 2019 human-like swine H3N2 spillover viruses. Further, SW/VA/19 was characterized in pigs in comparison to a contemporary clade 2010.1 isolate A/Swine/Missouri/A01410819/2014 (H3N1) (SW/MO/14) and to that of a human seasonal virus A/Switzerland/9715293/2013 (H3N2) (A/SWITZ/2013) in the same 3C3a clade as SW/VA/19. By studying this spillover through computational, in vitro, and in vivo experimental approaches, the risk for pig populations is better defined.

## 2. RESULTS

### 2.1 Genetic characterization of sequenced spillovers

To provide insight into the relationship of the four H3N2 spillover detections to human seasonal influenza, the genome sequence of SW/VA/19 was compared to that of human seasonal IAV (Fig. 1). All eight segments of SW/VA/19 matched closely to IAV from the 2018-2019 U.S. human influenza season with 99.7%-100% nucleotide identity. Within the HA and NA glycoprotein segments, little to no differences were detected in amino acids for SW/VA/19 versus one or more human IAVs sequenced from that flu season (Fig. 1A, B). There were minimal amino acid differences between viruses in the HA and NA segments of all four swine spillover detections (Supplemental Figures 1 and 2). This included no discernable amino acids differences in HA antigenic, receptor binding or HA1/HA2 cleavage sites. There were also no key differences in enzymatic active sites in the NA or in predicted glycosylation sites in either the HA and NA among the swine detections in comparison to the to the A/Kansas/14/2017 strain that was a component of the Northern Hemisphere 2019–2020 seasonal human influenza vaccine (18) The A/Kansas/14/2017 vaccine strain and the SW/VA/19 strain had only three amino acid differences in the HA at positions Q173K, M346L and A530V (Fig. S1). Collectively this analysis confirms little to no mutation was required to facilitate human-to-swine spillover with these viruses. Phylogenetic characterization of the HA segment including the four swine sequences revealed the two Virginia spillover detections were likely independent to that of the Illinois and Michigan spillover detections, as they were phylogenetically distinct (Fig. 1C).

**Figure 1.**
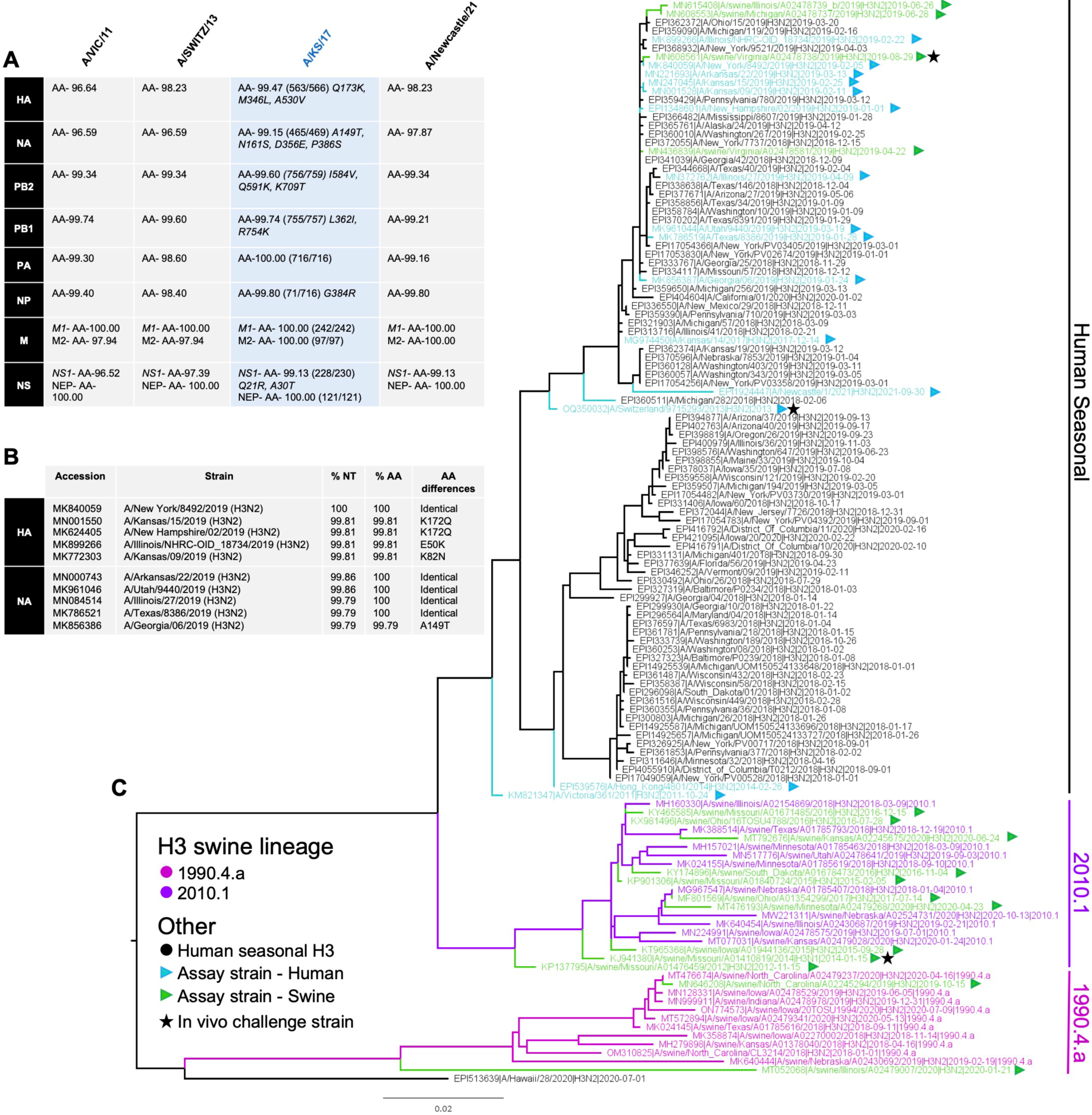
Genetic relationship of SW/VA/19 and human seasonal H3N2 viruses. **(A)** The amino acid (AA) identity of SW/VA/19 gene segments compared to North America seasonal H3N2 vaccine strains A/VIC/11, A/SWITZ/13 and A/KS/17-A/Kansas/17/2017 (shaded blue, closest seasonal vaccine match) and the first publicly accessible sequenced 3c3a1 clade virus detected in 2021 (A/Newcastle/1/2021) identified in NextStrain (https://nextstrain.org/flu/seasonal/h3n2/ha/2y accessed 2/11/2022) (19). **(B)** Percent nucleotide (NT) and amino acid (AA) identity of SW/VA/19 compared to the top 5 NCBI BLAST matches for HA and NA genes of human influenza. All amino acid position differences were calculated based on pairwise alignments using MUSCLE. **(C)** Maximum-likelihood tree showing the relationship of human seasonal H3 and swine H3 viruses detected between 2018 and 2020. Triangles denote strains collected from humans (blue) or swine (green) that were utilized in the genetic comparisons or in subsequent *in vitro* assays. Stars (black) denote three *in vivo* challenge strains.

### 2.2 Viral replication kinetics of human viruses, 2010.1 contemporary viruses, and SW/VA/19 in cell lines

To determine the extent SW/VA/19 replicates in different cell lines, a series of cell culture infections were performed comparing SW/VA/19 viral titers using a panel of 14 viruses as described in Table 1. The phylogenetic relationships of these viruses are shown in Figure 1C. The virus panel included eight clade 2010.1 swine H3 IAV strains (SW/MO/12, SW/MO/14, SW/MO/15, SW/IA/15, SW/MO/16, SW/SD/16, SW/OH/17, SW/MN-20), two clade 1990.4 swine H3 strains (SW/NC/19, - SW/IL/20) and three representative human IAV H3N2 strains of previous seasonal human vaccines (A/VIC/11, A/SWITZ/13, A/HK/14). Viral kinetics were assessed in swine-derived immortalized swine respiratory epithelial cells (iSRECs) (Fig. 2A-B) and in Madin Darby Canine Kidney (MDCK) cells at physiologically relevant basal body temperatures of 37 °C for human and 40 °C for swine (Fig. 2C-D). Viral kinetic data revealed the SW/VA/19 representative spillover strain (Fig. 2, orange line and circle) reached peak viral titers in iSRECs that were on average 1.5 to 4 log_10_ higher than that of the three human strains (Fig 2, black lines) at both 37 °C and 40 °C, but lower than all clade 2010.1 and 1990.4 swine viruses tested. The reduced replication of the human H3N2 viruses was attributed to both the iSREC cell line and the elevated temperature as these viruses amplified to >7 log_10_ on average in MDCK at 37°C (Fig. 2C) but significantly lower at 40 °C (Fig. 2D). A subset of the viruses in the panel were tested in primary normal human bronchial epithelial (NHBE) cells (Fig. 2E) and were consistent with MDCK findings, with the highest viral titers of >10^7 log_10_ observed with the human viruses A/VIC/11 and A/SWITZ/13 that were the most restricted viruses in swine cells.

**Figure 2.**
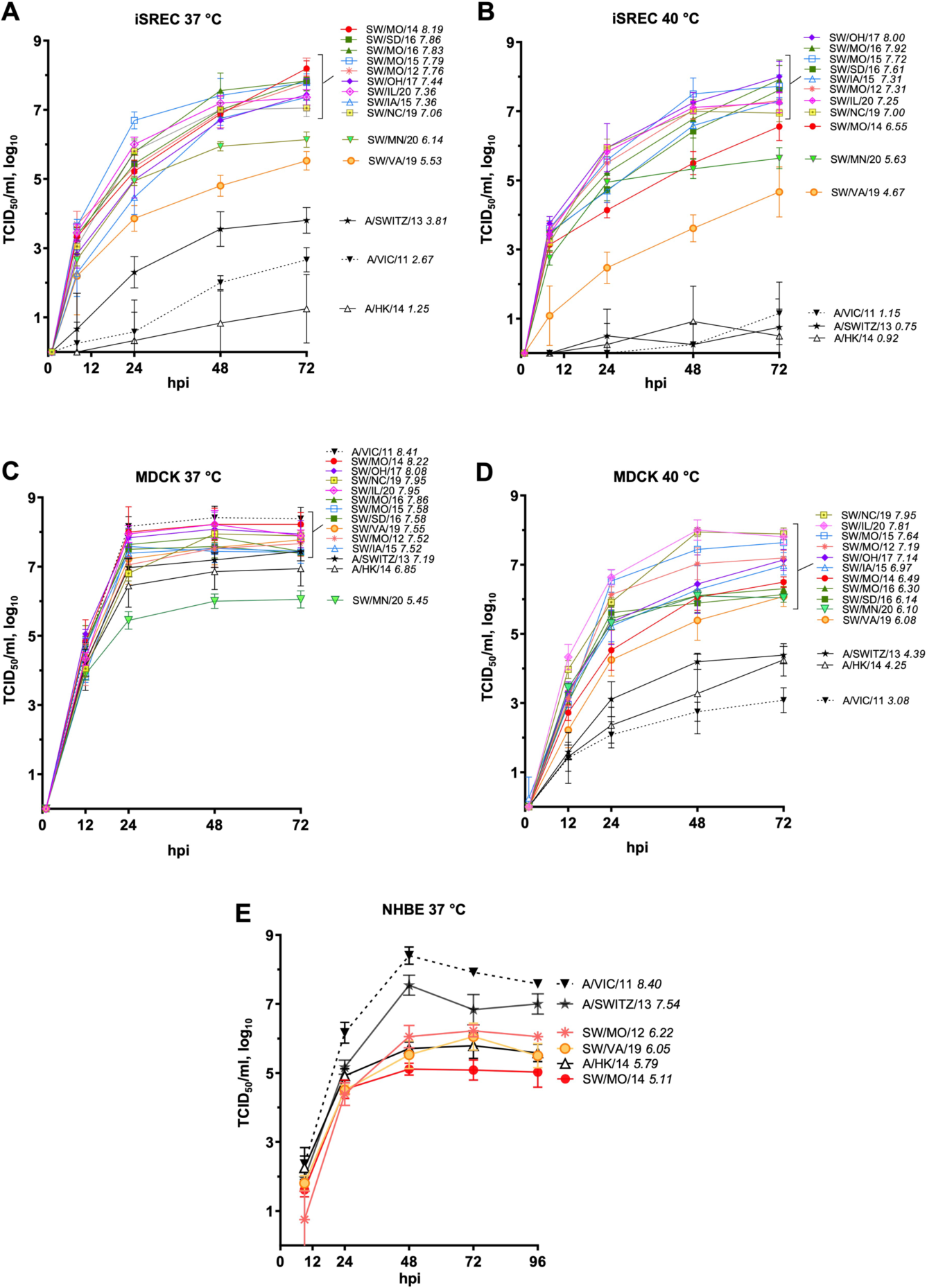
Viral titer kinetics of three human H3N2 viruses and eleven swine H3 viruses at 37 °C and 40 °C. A panel of viruses were amplified in iSREC **(A-B)**, MDCK **(C-D)**, and a subset of viruses in primary normal human bronchial epithelial (NHBE) cells **(E)**. Supernatant was analyzed by TCID_50_ with a readout by hemagglutination assay over 72- or 96-hours post infection (hpi). Data is from two independent experiments each performed in triplicate (n=6) with a starting MOI of 0.01. Data presented are group means ± standard error of the means. Human viruses are indicated by black; SW/VA/19 by orange; and swine viruses in other respective colors. Noted in figure legend in italics is the mean log_10_ maximum peak viral titer during the virus infection time course.

**Table 1.**
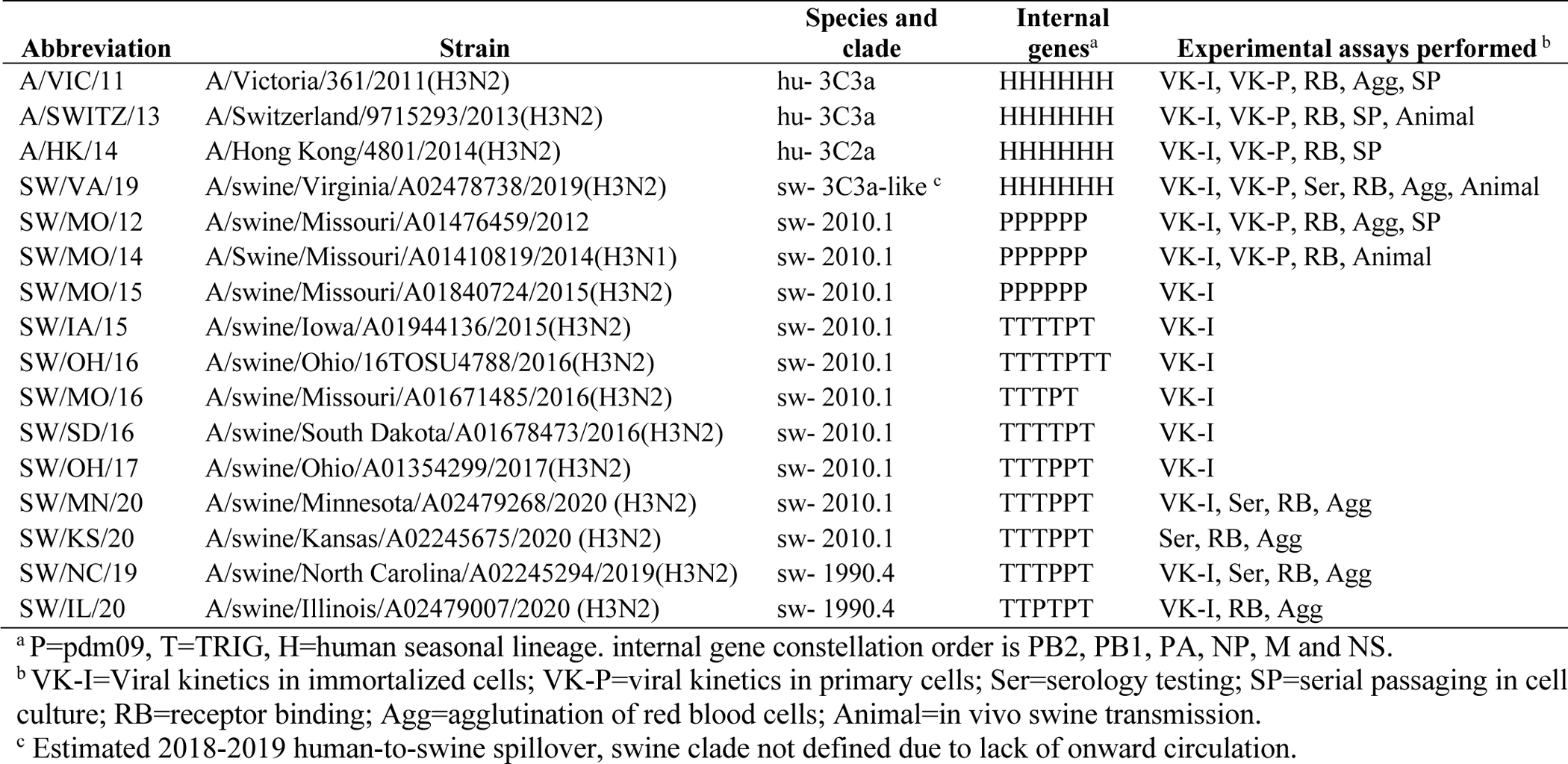
Viruses characterized and experimental assays performed.

To further define the replication properties of SW/VA/19 in swine using a subset of viruses shown in Figure 2, virus replication was characterized in primary swine epithelial cells isolated from two pigs. Primary swine cells extracted from digested nasal turbinates, trachea, and lung swine tissue were plated directly into 24-well tissue culture plates without cell passaging (passage zero). Cells from two distinct animals (Fig. 3) both confirmed SW/VA/19 replicated to intermediate levels, often higher that of the three human H3N2 viruses but lower than two representative swine viruses from the established 2010.1 lineage in both nasal turbinates and trachea cells (Fig. 3A-D). In lung swine cells (Fig. 3E-F), SW/VA/19 and the three human viruses replicated to comparable levels, but lower than that of the SW/MO/12 and SW/MO/14 2010.1 viruses. Immunofluorescence staining of primary cells showed similar α2-3 and α2-6 sialic acid (SA) profiles between both animals, with more pronounced α2-6 SA in cells isolated and cultured from the trachea and lower lung regions versus the nasal turbinate cells. Primary swine cell data demonstrated SW/VA/19 did not replicate in the three differing swine respiratory cell cultures as efficiently as the two 2010.1 contemporary swine viruses.

**Figure 3.**
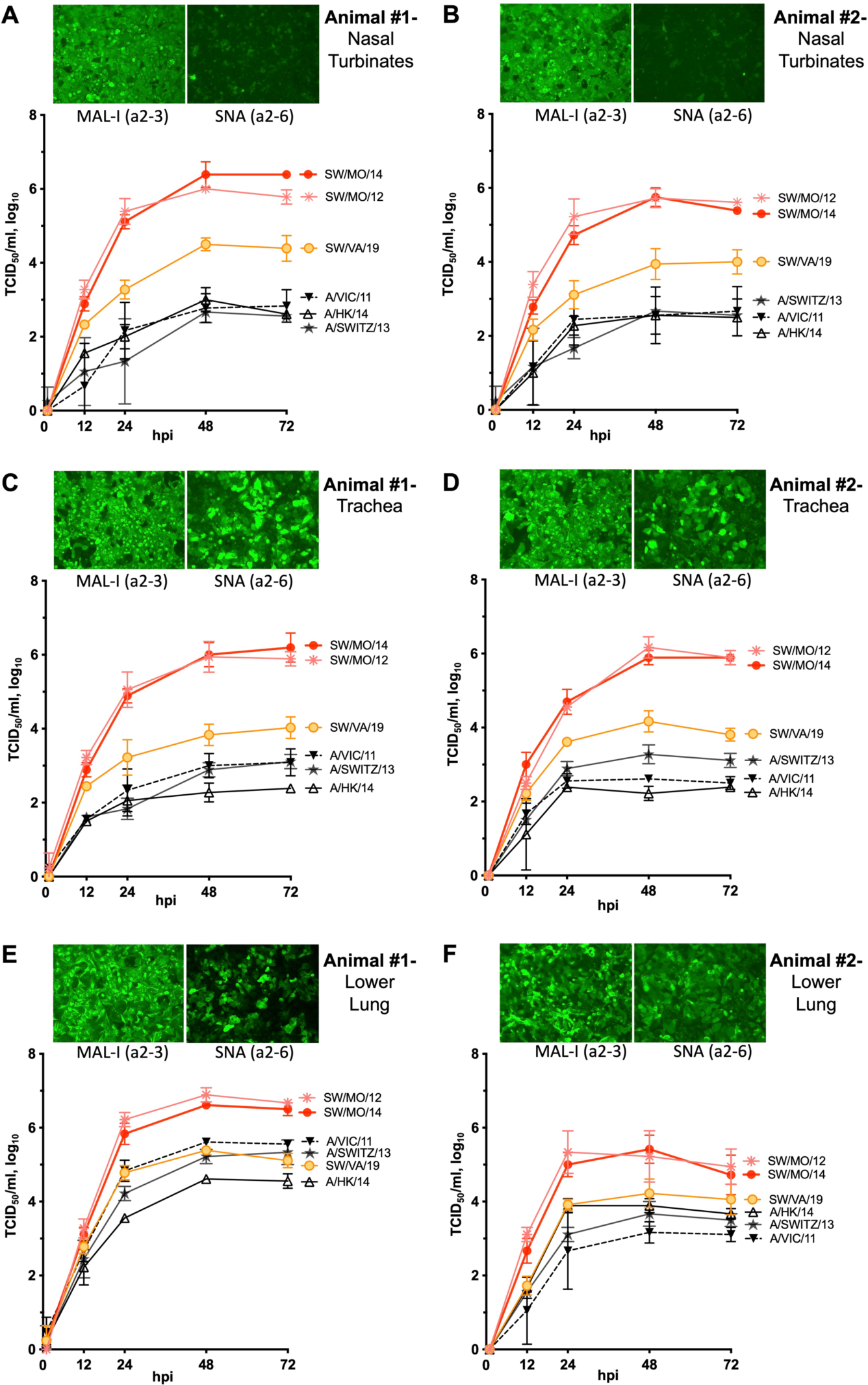
Viral titer kinetics in primary swine cells isolated from two different animals. Primary tissue from nasal turbinates **(A-B)**, trachea **(C-D)**, lung **(E-F)** were cultured and infected, then supernatant was analyzed by TCID_50_ with a readout by hemagglutinin assay at 1, 12, 24, 48, 72 hpi. Data is from two independent experiments each performed in triplicate (n=6) with a starting MOI-0.01. Data presented are group means ± standard error of the means. Human viruses are indicated in black; SW/VA/19 in orange; and swine viruses in pink and red. Representative immunofluorescence images from the animal tissue staining for biotinylated alpha 2-3 sialic acid (MAL-I lectin) or alpha 2-6 sialic acid expression (SNA lectin) as detected with Streptavidin AlexaFluor™-488 (green).

### 2.3 Survival of SW/VA/19 at intermediate titers when propagated in swine respiratory cells

Since SW/VA/19 replicated less efficiently in iSRECs (Fig. 2) and primary swine nasal turbinate and trachea cells (Fig. 3) compared to other swine viruses, we repeated propagation in swine iSRECs to assess fitness and observe potential changes. Viruses were passaged in 12 independent virus replicates for SW/VA/19 and the three human IAV’s A/VIC/11, A/SWITZ/13, and A/HK/14, with 6 independent virus replicates for SW/MO/12 and SW/MO/14 (Fig. 4A). None of the replicates of human viruses were recovered past passage 2 in iSRECs and were discontinued at passage 4 as they were qPCR IAV-negative. Only one SW/VA/19 isolate remained at passage four, and this isolate was expanded out to 24 additional replicates. No SW/VA/19 replicates were detected above 10^4 log_10_ viral titers, but 19 of 24 replicates from the expanded passage-4 isolate were recoverable at the end of passage 12.

**Figure 4.**
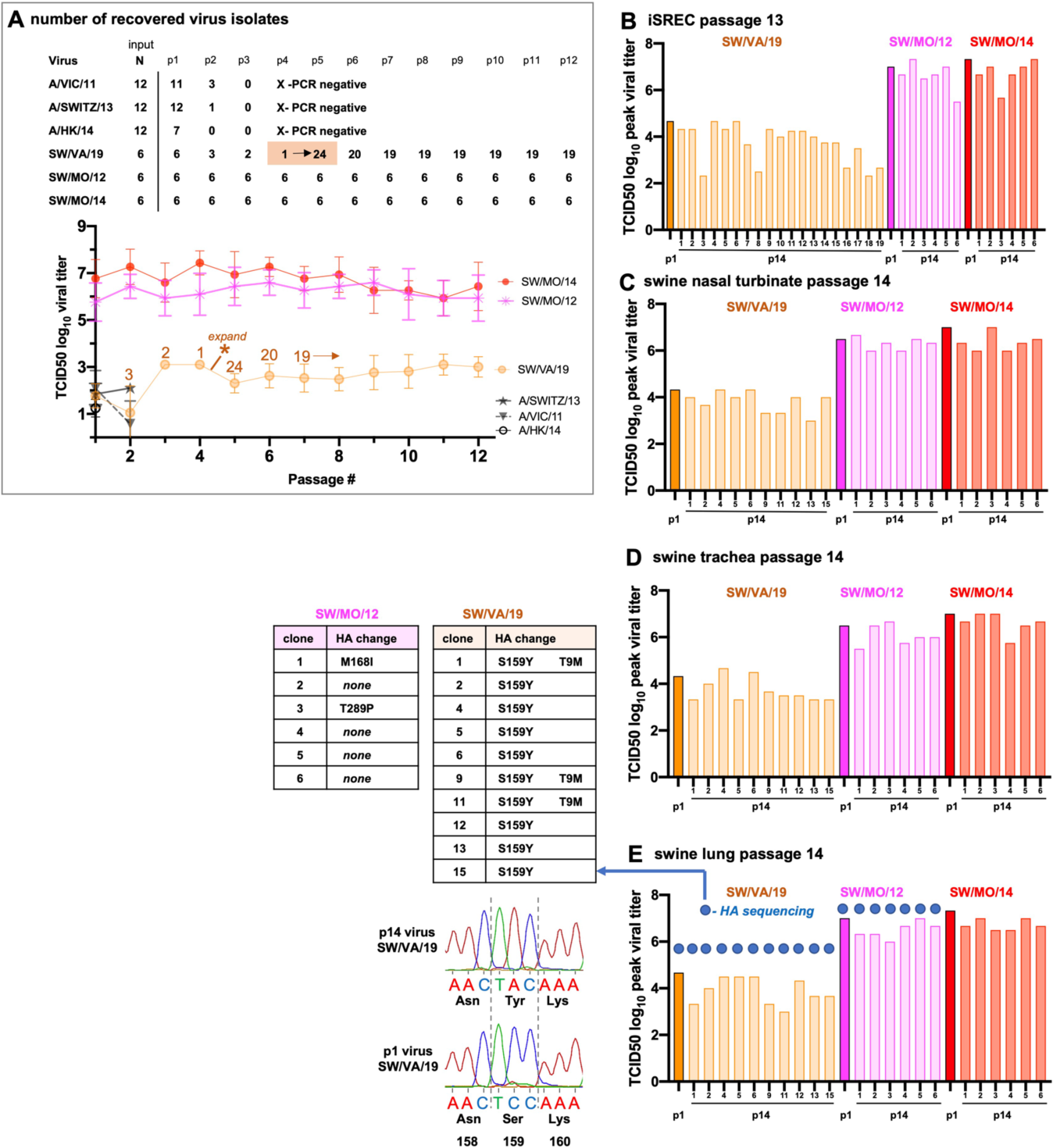
Survival during repeat propagation in swine cells. **(A)** Six viruses with n= 6 or 12 independent replicates each for A/VIC/11, A/SWITZ, A/HK/14, SW/VA/19, SW/MO/12 and SW/MO/14 were used to infect iSREC cells for 72 hr. Virus cell supernatant was 10-fold serially diluted and placed onto iSRECs with the infection process repeated for a total of 12 serial virus passages with viral titers monitored throughout. Human viruses were not isolated after passage 2. A single SW/VA/19 virus replicate remained at passage 4 and was expanded to 24 replicates (asterisks). **(B)** Virus at passage 12 from iSREC supernatants were used to infect iSRECs again (virus passage 13) at an MOI of 0.1 to compare with passage 1 stock virus (dark vs. light colored bars). The passage 13 viruses in panel B were used to infect each of the swine primary **(C)** nasal turbinate cells, **(D)** trachea cells, and **(E)** lung bronchial cells to represent passage 14 viruses. All peak viral titer bar graphs represent peak viral titers over a 96hr kinetic time point assessment. Viral titers are the mean of two independent samples (n=2). Blue circles in panel E denote samples subjected to HA gene sequencing and the amino differences between passage 1 virus and passage 14 virus are presented to the left of the graph.

The virus titers of all passage 12 virus isolates were compared to passage 1 stock virus in iSRECs using a starting input of MOI 0.1. The resulting passage 13 iSREC viral titers showed no enhanced replication (Fig. 4B). The passage 13 iSREC SW/VA/19 replicates with the 10 highest viral titers were next used to infect primary swine cells (passage 14). Higher viral replication was not observed in primary swine respiratory cells from either nasal turbinates, trachea, or lung (Fig. 4C-E). The passage 14 virus was subjected to sequencing of the viral HA segment after infecting swine lung cells (Fig. 4E, circles). Sequencing confirmed no virus cross-contamination had occurred and identified a serine to tyrosine HA change at position 159 (S159Y) in all 10 sequenced SW/VA/19 viruses, likely associated with the expansion of the single SW/VA/19 virus sample from passage 4 (Fig. 4A, asterisks). A Tyrosine to Methionine change at HA position 9 (T9M) was also identified in 3 of 10 viruses. HA sequencing of all six SW/MO/12 passaged viruses identified one virus containing a M168I change, a second virus with a T289I change and four viruses with no amino acid changes. Repeat propagation on iSREC cells demonstrated SW/VA/19 retained lower viral titers in swine cells compared to 2010.1 strains but higher than the human seasonal strains. A change occurred at an antigenically relevant site (position 159) that was independent of immune pressure. Additional attempts with MOI 1.0 and higher supernatant volume (higher 1:5 transfer inoculum) also proved unsuccessful in maintaining human A/VIC/11, A/SWITZ/13, A/HK/14 viruses in iSRECs past passage 3.

### 2.4 Antigenic and receptor binding characteristics of SW/VA/19

To further evaluate the relationship of SW/VA/19 with the two major 1990.4 and 2010.1 swine H3 clades that circulate in North America, serology and receptor binding assays were performed using a subset of Table 1 viruses (Fig. 5). The four 2019 spillover detections were similar to one another but had amino acid differences within key antigenic and receptor binding HA positions compared to currently circulating swine H3 clade 1990.4 and 2010.1 viruses (Fig. 5A, blue and pink). First, hemagglutination inhibition (HI) assays were performed using monovalent antisera raised in vaccinated pigs corresponding to SW/MN/20 and SW/KS/20 from 2010.1 clade, and SW/NC/19 from 1990.4 clade. These three virus strains were selected as clade consensus representatives (17). HI data demonstrated SW/VA/19 had at least an 8-fold loss in titer to the heterologous swine IAV anti-sera (Fig. 5B), indicating a significant loss of cross-reactivity between SW/VA/19 and the 1990.4 and 2010.1 swine viruses.

**Figure 5.**
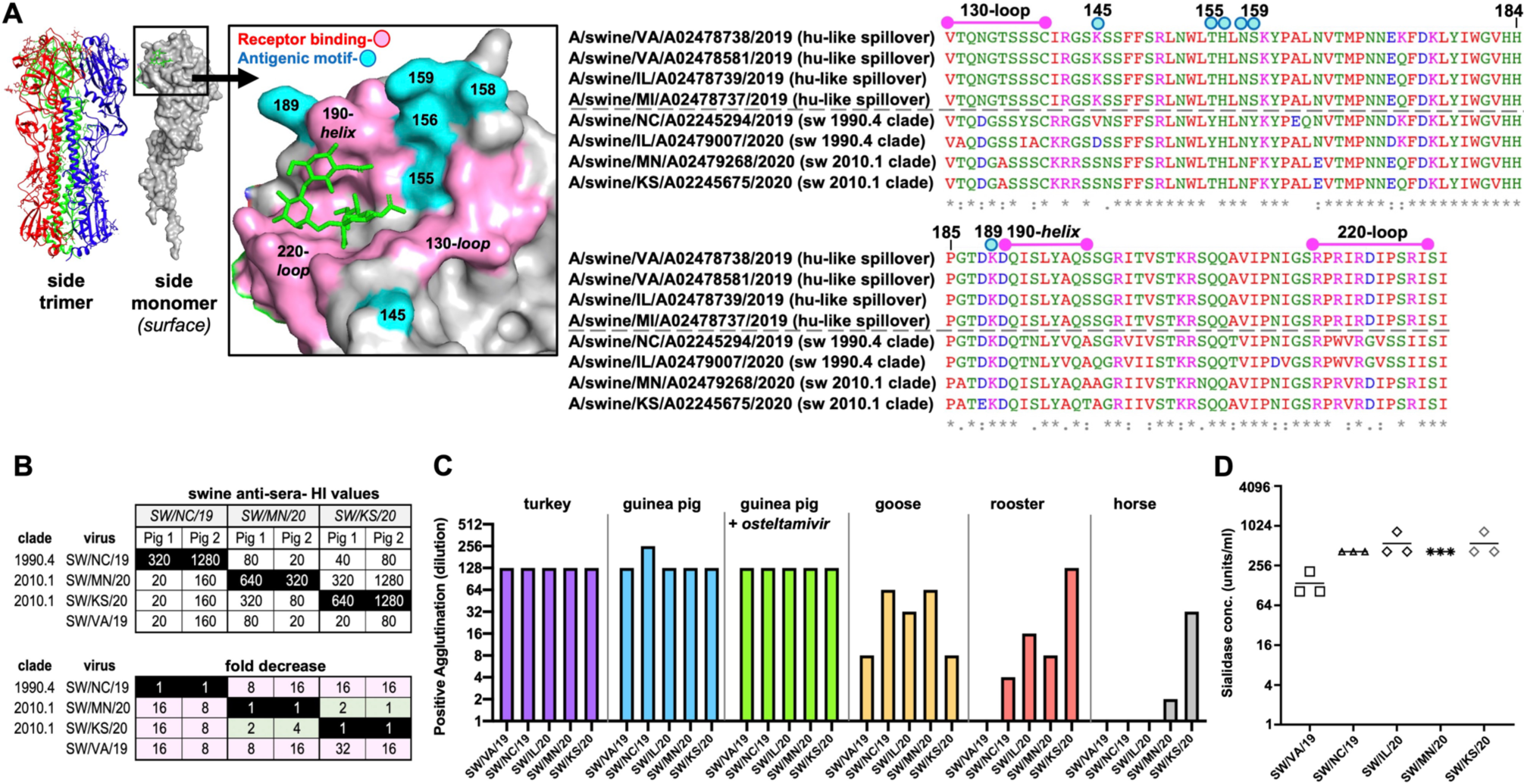
Antigenic and receptor binding relationship of the 2019 spillover virus detections compared to clade 1990.4 and 2010.1 swine H3N2 viruses. **(A)** The HA receptor structure shown in trimer and monomer form. Adapted from the solved A/Michigan/15/2014 (H3N2) hemagglutinin in complex with 6’-SLN (green) (https://doi.org/10.2210/pdb6bkt/pdb) (22). Key antigenic sites at positions 145, 155, 156, 158, 159, and 189 are in cyan. Three HA domain regions associated with receptor binding are in pink (23). HA alignments for the four 2019 swine spillover detections and the two representative strains from the contemporary 1990.4 and 2010.1 swine H3 clades that circulate in North America. **(B)** Hemagglutination inhibition (HI) titers and fold-change from homologous with a panel of monovalent antisera produced in vaccinated swine. Homologous titers from antigen paired antisera are highlighted in black with corresponding fold-change reductions below. **(C)** Viruses were assayed for agglutination of red blood cells (RBCs) by standard hemagglutination (HA) assay from turkey, guinea pig (without and with Oseltamivir; Ost), goose, rooster, and horse. **(D)** Concentration of sialidase treatment of turkey RBCs for 1 hour resulting in positive agglutination (virus at 8 HAU/50 μl) done in triplicate to quantify the relative receptor binding affinity.

Next, the receptor-binding properties of the four viruses tested by serology and an additional 1990.4 virus SW/IL/20 (Table 1) were assayed for ability to bind and agglutinate fresh red blood cells (RBCs) isolated from different species (turkey, rooster, guinea pig, goose, and horse) by standard hemagglutination (HA) assay (Fig. 5B). All viruses efficiently agglutinated turkey and guinea pig RBCs, which express SAα2-3Gal and SAα2-6Gal receptors on their cell surfaces. Virus binding was also minimally impacted in the presence of 10 μM oseltamivir, a compound that binds in the active site of NA (20). SW/VA/19 agglutination efficiency was lower in goose and rooster RBCs, a trend also observed in the other swine viruses. Horse RBCs that display mainly SAα2-3Gal (16) were minimally agglutinated apart from moderate binding by SW/KS/20. Some human H3N2’s evolved to inefficiently agglutinate erythrocytes (21), but this data demonstrates SW/VA/19 agglutinated turkey and guinea pig RBCs, which are critical for antigenic characterization.

Erythrocyte-virus associations were further profiled by treating turkey RBCs with differing concentrations of Clostridium perfringens neuraminidase (sialidase) that remove α2-3, α2-6, and α2-8 linked SA as a measure of relative virus binding affinity (Fig. 5D). The four 1990.4 and 2010.1 swine viruses had similar binding profiles that trended higher than that of SW/VA/19, indicating HA binding affinity was diminished for SW/VA/19 compared to swine viruses.

### 2.5 Infection, transmission, and pathogenesis of SW/VA/19 in pigs

Ten pigs in each of three biocontainment rooms were challenged with either SW/VA/19, SW/MO/14 clade 2010.1 (13), or human seasonal A/SWITZ/13 virus to compare shedding and transmission phenotypes. Five additional indirect contact pigs were placed in each room two days after initial infection in a pen physically separated by a 2-foot space. Nasal swab titers from challenged pigs (Fig. 6A) and contact pigs (Fig. 6B) demonstrated differences in kinetics as well as magnitude of titers. In primary challenged pigs, SW/MO/14 was shed in 9 of 10 pigs at 1 day post infection (dpi) with a peak group mean viral titer of 3.7 log_10_ TCID_50_/mL. SW/VA/19 virus was detected in 7 of 10 pigs and A/SWITZ/13 in 5 of 10 pigs at 1 dpi. On 2 dpi, 9 of 10 SW/VA/19 pigs and 10 of 10 A/SWITZ/13 pigs shed virus. By 5 dpi, SW/VA/19 was shedding significantly higher titers on average (4.1 log_10_ TCID_50_/mL) than SW/MO/14 (1.8 log_10_ TCID_50_/mL). SW/MO/14 and SW/VA/19 viruses displayed similar peak titers in pigs, but SW/VA/19 virus nasal shedding was delayed.

**Figure 6.**
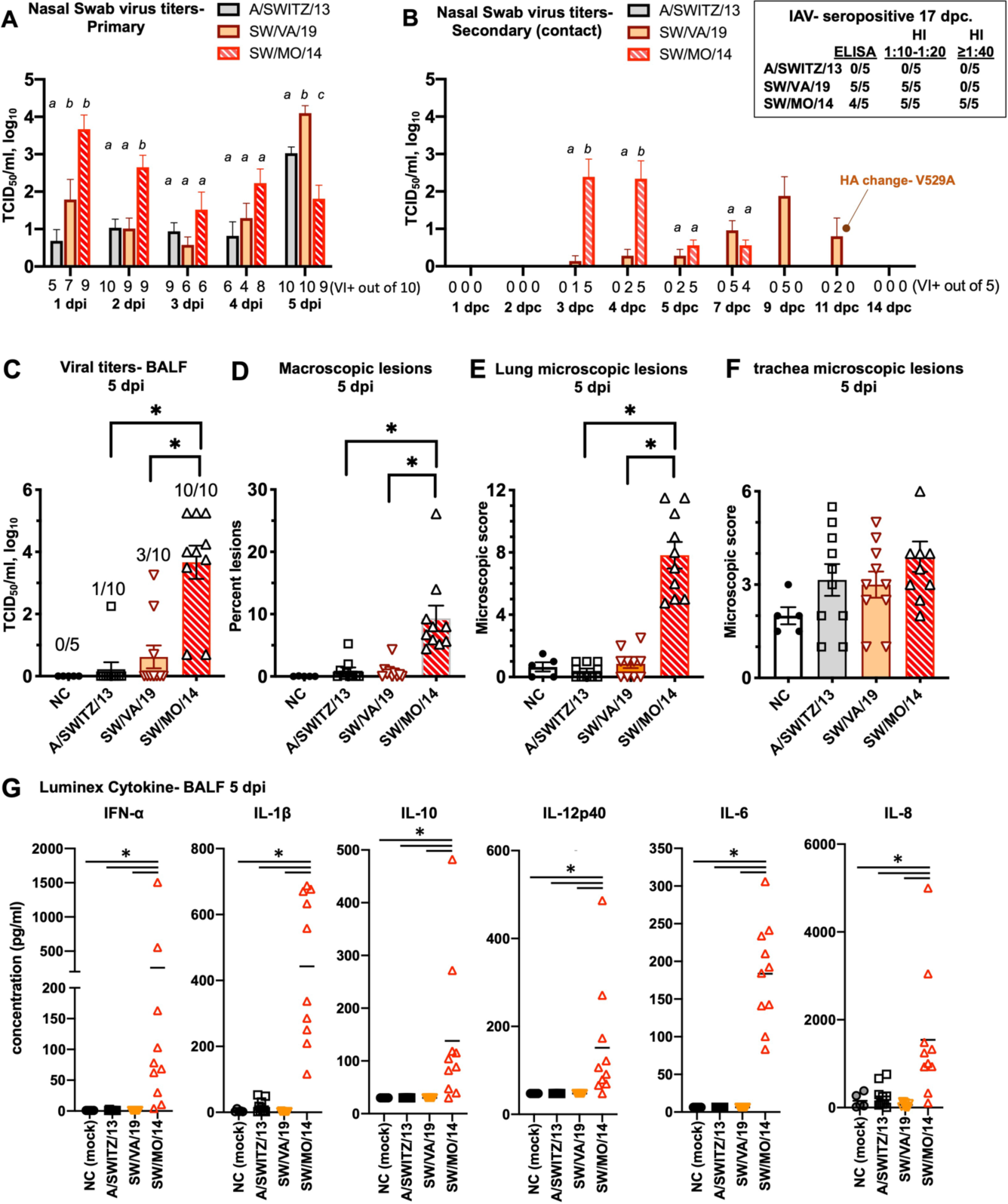
Infection, transmission, and lung lesions in pigs. Nasal swab samples were collected every day in **(A)** challenged pigs (n=10) or **(B)** indirect contact pigs (n=5) and titrated in tissue culture. Treatment group means with statistically significant differences (P ≤ 0.05) are identified by different lowercase letters above the group mean bars (a, b, c). The number of pigs positive by virus isolation (VI) are noted below the X-axis. The HA of two SW/VA/19 positive pigs at 11 days post contact (dpc) were sequenced, revealing a single Valine-to-Alanine HA change (V529A). At 17 dpc sera from contact pigs were analyzed by ELISA and hemagglutination inhibition (HI) assay for IAV-specific antibodies (inset box). **(C)** The viral titers and the number of challenged pigs in which virus was recovered from BALF samples. **(D)** The percentage of macroscopic lung lesions over the surface area of the extracted lung was measured as a correlate of lung damage inflicted by the virus. **(E)** Histopathology composite lesion scores from examined lung based on six parameters and from **(F)** Trachea sections based on two parameters as a measure of disease severity **(G)** Luminex porcine-specific cytokine levels in BALF samples were measured as a correlate of infection host response within the lung. All data presented are group means ± standard error of the means for all pigs in each respective group. Asterisks in (C) and (G) denote statistically significant differences (P ≤ 0.05) in virus groups compared to SW/MO/14.

SW/VA/19 was detected in nasal swabs from all five contact pigs by 7 days post contact (dpc), whereas all five contact pigs in the SW/MO/14 group were positive by 3 dpc (Fig. 6C). No virus was recovered from contact pigs for the A/SWITZ/13 group. Sera analyzed at 17 dpc in contact pigs confirmed viral transmission data as IAV-specific antibodies were detected by ELISA and HI assay methods (Fig. 6B, right inset). The delay in transmission to contact pigs was reflected in low HI titers, but seroconversion was detected in all 5 contact pigs. In the 2 pigs shedding SW/VA/19 virus at 11 dpc, the segment encoding the HA gene was sequenced. Compared to input challenge virus, a single Valine-to-Alanine amino acid change at position 529 in the HA was identified in both animals.

Next, an assessment into how the three viruses used in animal studies impacted the lungs of challenged pigs was examined. Viral titers from bronchoalveolar lavage fluid (BALF) at 5 dpi were analyzed and on average SW/VA/19 (0.6 log_10_ TCID_50_/mL) and A/SWITZ/13 (0.2 log_10_ TCID_50_/mL) groups had significantly lower mean viral titers compared to SW/MO/14 (3.7 log_10_ TCID_50_/mL) (Fig. 6C), with virus recovered in 3 of 10 SW/VA/19 and 1 of 10 A/SWITZ/13 pigs. SW/MO/14 induced significantly higher percentages of macroscopic lung lesions on average (9.3%) compared to SW/VA/19 (0.7%) and A/SWITZ/13 (0.9%) (Fig. 6D). Histopathological examination of lung sections for microscopic lesions (Fig. 6E) agreed with macroscopic findings, but no significant differences in pathologic changes were observed between viruses in the trachea (Fig. 6F). Host cytokine levels (IFN-α, IL-1β, IL-10, IL-12p40, IL-6 and IL-8) in BALF for SW/MO/14 were indicative of viral infection for SW/MO/14, while little to no cytokine induction for SW/VA/19 and A/SWITZ/13 was evident as compared to that of nonchallenged (NC) BALF samples (Fig. 6G). Figure 6 demonstrates SW/VA/19 was transmissible, albeit at a slower rate than SW/MO/14 but resulted in minimal disease in experimentally challenged pigs.

## 3. DISCUSSION

The first detection of a spillover from the 2018-2019 H3N2 human influenza season was from a pig respiratory specimen submitted to the Iowa State University Veterinary Diagnostic Laboratory in April 2019. Three additional strains were detected over the next six months. These detections represent at least two independent human-to-swine introductions based on phylogenetic analyses. Epidemiologic links between the four detections were not known. The extent that the 2019 human spillovers could transmit in pigs was unclear as these strains were fully human-derived viruses with little genetic differences to that of seasonal human H3N2 influenza from the 2018-2019 flu season and there was limited evidence of onward spread beyond the index herds. As the barriers that prevent fully-human H3N2 viruses to spread through the pig populations are complex and not fully understood (24), we demonstrated that reassortment was not required for these human seasonal H3N2 to transmit in pigs.

IAVs have moderately high mutation rates and constantly evolve to adapt to changes in the host environment and to transmit across species. The adaptation of IAV in pigs is largely attributed to reassortment and point mutations (24). As this study demonstrates, the human-like internal gene constellation (HHHHHH) with high similarity to human seasonal H3N2 remained intact and was transmissible in pigs. These data suggest the species barriers for future introductions of human H3N2 might be lower than previously expected. Virus was detected in nasal secretions over multiples days and at similar levels as SW/MO/14 in experimentally infected pigs; however, SW/VA/19 was less efficient in viral replication and avirulent in pigs, with delayed transmissibility and minimal replication in the swine lung. Since infection of SW/VA/19 led to asymptomatic outcomes in pigs, its unknown if additional infections went unreported in the field. No additional detections of these viruses or remnants of human seasonal virus “H” internal gene segments through reassortment with endemic swine viruses were reported by USDA surveillance sequencing efforts, thus, these viruses did not likely spread in U.S. pig populations. If these spillovers were to be sustained, they could potentially complicate vaccine efforts since there was diminished serum cross-reactivity for spillover virus against the two major lineages of H3N2 that co-circulate in U.S. swine that are currently the targets of vaccine efforts in swine populations.

An important question that remained unaddressed is what genetic changes allowed the SW/VA/19 progenitor to cross from humans into pigs that did not otherwise happen in other human influenza seasons? Prior work demonstrated that A/Victoria/361/2011 (A/VIC/11), did not transmit to contact pigs and challenged pigs had minimal shedding (3 dpi only) (13) and our findings are in agreement with that study. Data presented here for A/SWITZ/13 from the same human seasonal H3 clade 3C3a did not transmit to indirect contacts, but unlike A/VIC/11, was shed by all primary pigs over multiple days. The combination of genetic differences between A/VIC/11 and A/SWITZ/13 that may confer increased nasal shedding is unknown at this time as amino acid differences spanned across all eight segments.

There are few publications assessing the genetic determinants for human-origin viruses to become swine-adapted. One recent study by Mo et. al (25) using viruses similar to our study (A/VIC/11 and SW/MO/14) offers some intriguing insight. Specifically, a reverse genetics A/VIC/11 virus with the original human seasonal HA and NA intact but encoding a pandemic-associated TRIG internal cassette (VIC11pTRIG virus) was used in animal transmission studies to overcome the limited transmission of the parent A/VIC/11 virus. These studies with VIC11pTRIG assessed virus changes through sequencing of nasal swabs from contact pigs. Variants with HA changes in respiratory contact pigs after transmission at either A138S, V186G, and F193Y were identified. The three HA positions reside within antigenic site B near or within the RBS. The three variants also showed more efficient replication in primary swine epithelial cells at most time points, implicating that receptor binding changes help facilitate human-to-swine adaptation. Two additional earlier studies have also implicated that the number of punitive glycosylation sites in the human H3 HA is a determinant for human-to-swine adaptation (16, 26). Phylogenetic data (26) that was later supported by experimental data (16) showed an increased antiviral neutralization by recombinant porcine surfactant D against A/VIC/11 versus A/SW/MO/14 that lost glycosylation of N133 and N165. In comparison to these studies, the only HA variant we detected in contact animals for SW/VA/19 was at HA position V529A that is not associated with RBS or glycosylation. SW/VA/19 propagated in swine cells resulted in variants of S159Y with or without T9M that is not associated with glycan sites. S159Y resides within the antigenic B-site and has RBS associations (27) but this change did not change swine viral titers in either primary or immortalized swine cells. The agglutination profiles and receptor binding affinity assays demonstrated lower binding affinity for SW/VA/19 compared to the dominant swine H3 viruses, suggesting a potential rationale for limited transmission and lower lung virus detection. Additional genetic factors, including those yet to be identified within internal genes segments (HHHHHH) for SW/VA/19 also likely played a key role in limiting transmission for SW/VA/19-like viruses in swine.

The continued threat of human H3N2 viruses to pigs is unknown. Global circulation patterns of influenza viruses were disrupted from social measures implemented against the SARS-CoV-2 coronavirus. During 2020-2022 there was a 99% reduction in detections in human influenza cases (28). The lack of detection of human H3N2-3C3a viruses was apparent due to only 3 virus sequences deposited into the GISAID global database during this timeframe with none from North America. In the most recent 2022-2023 flu season there continued to be a lack of 3C-clade detections globally, with the predominant detection being subclade 3a.2. Despite the prolonged absence of influenza viruses of 3C3a, continual swine surveillance is paramount. A better understanding of the propensity for spillover at the human-swine interface allows for better informed policy decisions tied to future vaccination strategies. Understanding the propensity for human influenza to spill into pigs also better addresses potential reservoirs for influenza evolution that could lead to future pandemics. In summary, this study demonstrates that human-to-swine spillover viruses descending from the human H3 3C3a clade shed and transmitted from challenged pigs. Although SW/VA/19 with an entirely human influenza genome was capable of infection and transmission in pigs, further mutation and reassortment may be necessary for sustained pig to pig transmission since endemic spread through the swine population was not yet observed.

## 4 MATERIALS AND METHODS

### 4.1 Viruses

Table 1 describes the 15 IAV strains used as part of this study with the phylogenetic relationship to one another highlighted in Figure 1C. Swine IAV isolates were obtained from the United States Department of Agriculture (USDA) Influenza A Virus in Swine Surveillance System (denoted by a nine-digit alphanumeric barcode beginning with A0 in the strain name). IAV sequences described throughout were obtained from the National Center of Biotechnology Information (NCBI) Influenza Virus Resource (IVR) (29) and the Influenza Research Database (IRD) (30). Additional information for the eight genome segments of IAVs used in pigs with the following accession numbers were downloaded from NCBI GenBank: SW/VA/19 [MN608558-MN608565]; SW/MO/14 [KJ941380-KJ941382, KR092079-KR092083]; or through GISAID: A/SWITZ/13 [EPI_ISL_166310]. To assess the diversity of the strains characterized in this study, all human H3 HA sequences collected in the United States between 2018 and 2020 (n=8,885) were downloaded from GISAID (31). All swine H3 HA sequences from the same timeframe (n=1,178) were downloaded from BV-BRC (Bacterial and Viral Bioinformatics Resource Center) and classified using octoFLU (32). Swine H3 sequences that were classified as belonging to the 1990.4.a (n=521) or 2010.1 (n=509) clades were kept, and the remainder were not used in subsequent analyses (n=148). The human seasonal H3, swine H3 1990.4.a, and swine H3 2010.1 lineage datasets were aligned separately using mafft v7.490 (33) and a maximum-likelihood tree was inferred for each alignment following automatic model selection with IQ-Tree2 v2.2.2.6 (34, 35). To representatively subsample the datasets, we used PARNAS v0.1.4 (36) with a defined number of sequences (n=75 human H3, n=10 swine H3 1990.4.a, n = 10 swine H3 2010.1). The selected sequences from these three datasets were combined with the strains used in the biological assays into one FASTA file. The final FASTA file was used to align the sequences and a maximum-likelihood tree was inferred from the alignment using the same methods as above. All viruses were propagated in Madin–Darby Canine Kidney (MDCK) cells (FR-58, London Line) to generate viral stocks with the HA and NA segments sequenced before and after challenge of pigs to confirm their identity.

### 4.2 Cell lines and viral titers

Swine immortalized respiratory epithelial cells (iSRECs) obtained from the University of Minnesota under a material transfer agreement were isolated and cultured in the growth conditions descried previously (37, 38). MDCK cells (FR-58, London line), were obtained from the CDC-Influenza Research Repository (IRR) and cultured as specified. For viral titer assessment, cells were cultured in 24-well plates to ∼95% confluency and then infected in triplicate wells for each time point in Opti-MEM™ medium (Thermo Fischer Scientific, Waltham, MA) supplemented with 1 µg/ml L-1-Tosylamide-2-phenylethyl chloromethyl ketone (TPCK, Worthington Biochemicals, Lakewood, NJ). The starting multiplicity of infection (MOI) was 0.01 with supernatant collected over 72 hours post-infection (hpi). Viral titers from cell culture supernatants were determined in triplicate from two independent experiments (n=6 total) in MDCK cells using the 50% tissue culture infective dose (TCID50 per milliliter) according to the method of Reed and Muench (39). Virus positive wells were determined through use of a HA assay using 0.5% turkey red blood cells.

### 4.3 Primary cell characterization

Normal human bronchial/tracheal epithelial cells (NHBEs) (Lonza Inc. Walkersville, NC) were cultured in BronchiaLife™ Basal Medium with provided supplements (LifeLine Cell Medium, Frederick, MD) for one additional passage before infection. For primary swine cell virus kinetics, tissue sections from nasal turbinates, trachea, and lower lung from two separate freshly euthanized pigs were extracted and digested overnight in a serum-free DMEM Pronase/DNaseI solution (Sigma-Aldrich, St. Louis, MO) using a protocol as described prior (40). Next, tissues were gently scraped with a sterile scalpel with dislodged cells passed through a Falcon^®^ Nylon 40μm cell strainer (Thermo Fisher Scientific, Waltham, MA). Fetal bovine serum (FBS) at 10% final concentration was used to neutralize protease, cells were pelleted, resuspended in BronchiaLife™ medium and placed in T-150 sized tissue culture treated flasks. Sixty minutes later unattached cells in the supernatant were gently transferred to Corning^®^ BioCoat™ Collagen I-coated 24-well plates (Thermo Fischer Scientific, Waltham, MA) using 2×10^5 cells/well. Medium was replaced the next day with lung differentiation medium (LifeLine Cell Medium, Frederick, MD) that contained retinoic acid for an additional 24 hours before performing viral kinetics in Opti-MEM with 0.1 µg/ml TPCK.

For confocal immunofluorescence imaging, swine primary cells at 80-90% confluency were fixed, permeabilized, blocked, and washed as specified using the Image-iT^®^ Fixation Permeabilization Kit (Thermo Fischer Scientific, Waltham, MA). Primary antibodies (Vector Laboratories, Newark CA) at 1:100 dilution were biotinylated Sambucus nigra lectin (SNA) and biotinylated Maackia Amurensis Lectin I (MAL-I) each incubated on cells at room temperature with gentle agitation for 4 hours. Secondary labelling was with Streptavidin, Alexa Fluor™ 488 Conjugate (Thermo Fischer Scientific, Waltham, MA) at 1:5000 dilution for 2 hours. After PBS washes, cells were mounted in ProLong™ Gold Antifade Mountant with DAPI overnight (Thermo Fischer Scientific, Waltham, MA) without permeabilizing the cells. Image acquisition at 20x magnification was acquired with identical exposure and laser intensity. Images were acquired with a Leica DM IRBE microscope with a Leica DFC500 digital camera and the Leica Application Suite version 3.7.0 (Leica Microsystems, Wetzlar, Germany) at 200x magnification.

### 4.4 Virus propagation in swine cells

Stock viruses at passage 1 were 10-fold serially diluted onto 96 well plates containing 90-95% confluent iSREC cells with infections progressing for 72 hours. Next, the most dilute HA assay positive well was then 10-fold serially diluted again in the 96-well format using 10 μl of supernatant (1:10 total volume). The infection process was repeated for a total of 12 serial virus passages every 72-96 hpi over a 2-month duration with viral titers monitored throughout. Viral titers were monitored by TCID50 HA and confirmed for cytopathological effects (CPE) by light microscope. At passage 13, the starting MOI was adjusted to 0.01 and iSRECs infected. Afterwards, the infected supernatant was again set to MOI 0.1 and used to infected primary swine lung cells (passage 14). For viruses that failed to be passage, sample from the next passage were confirmed to be IAV-negative by real-time qPCR that targets a highly conserved M-gene segment region according to the manufacturer instructions (VetMax™ Swine Influenza Kit, Thermo Fischer Scientific, Waltham, MA). cDNA generated from total RNA from infected cell lines subjected to sanger sequencing confirmed no mixed-infections or cross-contamination had occurred from passaging after inspection of electropherograms of full-length HA sequences.

### 4.5 Serology and antigenic characterization

Swine antisera previously produced by immunizing each of two pigs was treated with receptor-destroying enzyme II (Hardy Diagnostics, Santa Maria, CA), heat inactivated at 56 °C, treated with 20% Kaolin (Sigma Aldrich, St. Louis, MO), and then absorbed with 50 % turkey red blood cells (RBC) to remove nonspecific hemagglutination inhibitors. HI assays were performed using wild-type viruses measured against 2-fold serial diluted treated serum with 0.5 % turkey RBCs with standard technique.

### 4.6 Agglutination and virus binding assays

For agglutination profiling, freshly isolated turkey, guinea pig, goose, rooster, and horse red blood cells (RBCs) were washed 3x and resuspended in ice cold PBS at a 1% vol/vol concentration. Viruses were then diluted to a concentration of 128-HAU/50 μl using turkey RBCs for read out. Next, virus at 128 HAU was then serial diluted 2-fold on 96-well V-bottom plates using cold PBS. All wells were tested against the four differing RBC types in addition to 1% guinea pig blood tested in the presence of 10 μM oseltamivir. Finally, equal 50 μl volumes of virus and RBC were incubated for 1hour and the highest dilution allowing RBC agglutination recorded. Each virus was assayed in duplicate and the mean value shown. The experiment was later repeated using differing batches of freshly collected RBCs. The mean value of the experiment performed in duplicate on two differing RBC extraction dates (n-4).

For measuring receptor binding avidity, sialidase (α2-3,6,8 Neuraminidase, New England BioLabs Inc. Ipswitch, MA.) was diluted 2-fold in PBS. Next, diluted sialidase was incubated in 10% turkey RBCs at 37 °C for 1 hour with inversion every 15 min. Sialidase was removed from RBCs through a series of 3 x 200 x g centrifugation steps with PBS buffer exchanges. Treated RBCs were then adjusted to a final 1% vol./vol. in PBS. Next, virus was incubated with 50 μl each of the sialidase-treated RBC samples for 1 hour in a 96-well plates at a final 8-HAU/50 μl concentration. Assays were performed in the 0 to 6656 sialidase units/ml final concentration range. The maximum sialidase concentration allowing for agglutination was recorded. Values reflects the assay performed independently three times using differing turkey RBCs and sialidase enzyme commercial lot numbers.

### 4.7 Animal study design

All pigs were cared for in compliance with the Institutional Animal Care and Use Committee of the USDA National Animal Disease Center. Pigs were treated with ceftiofur crystalline free acid and tulathromycin (Zoetis Animal Health, Florham Park, NJ) and were seronegative to IAV antibodies by a commercial ELISA kit (Swine Influenza Virus Ab Test, IDEXX) prior to the start of the study. At 4 to 5 weeks of age (0 days post-infection; dpi), 3 ml of 1 × 10^5^ TCID50/ml of either A/SWITZ/13, SW/MO/14, or SW/VA/19 challenge virus (Table 1) were delivered to pigs intratracheally (2 ml) and intranasally (1 ml). Pigs were challenged under anesthesia, using an intramuscular injection of ketamine (8 mg/kg of body weight; Phoenix, St. Joseph, MO), xylazine (4 mg/kg; Lloyd Inc., Shenandoah, IA), and Telazol (6 mg/kg; Zoetis Animal Health, Florham Park, NJ) cocktail. Ten primary pigs were challenged with each virus, while five pigs were not challenged (NC) as negative controls. Nasal swabs were collected daily before challenge (0 dpi) until 5 dpi, with virus isolation as previously described (41). At two dpi, five contact pigs for each respective group were placed in a separate raised decks in the same containment room. Contact pigs were in a pen separated from challenged pigs by an approximate 2 foot alley to assess indirect transmission dynamics. Contact pigs were swabbed daily 0 to 5 days post contact (dpc) and every other day thereafter until 14 dpc. Primary pigs at 5 dpi and contact pigs at 17 dpc were humanely euthanized with a lethal dose of pentobarbital (Fatal Plus, Vortech Pharmaceuticals, Dearborn, MI). BALF were collected in minimal essential media (MEM) from primary pigs at 5 dpi for analysis. ELISA (IDEXX) and HI assays were performed on serum from contact pigs at 17 dpc to determine if IAV-specific antibodies were present to indicate virus transmission.

### 4.8 Pathological examination and virus detection

At 5 dpi necropsy, the percent of lung surface affected with pneumonia was calculated as previously described (42–44). Lung microscopic lesions were blindly scored according to previously described parameters (44) and individual composite scores for each pig were computed, with a maximum composite score ranging from 0 to 22 for lung and 0 to 8 for trachea. Virus isolation (VI)-positive nasal swabs and BALF were titrated in MDCK cells as previously described (41, 45) and TCID_50_/ml virus titers were calculated for each sample by the Reed and Muench method (46). The VI detection limit was 10^0.7 (200 μl sample). Unrecoverable virus below the VI limit was recorded as a value of zero for statistical purposes.

### 4.9 Cytokine analysis

Luminex^®^ was performed on 25 μl of BALF sample from all virus challenge and mock pigs through use of the Cytokine & Chemokine 9-Plex Porcine ProcartaPlex™ 2.5 kit (Thermo Fischer Scientific, Waltham, MA) as instructed.

### 4.10 Statistical analysis and data availability

Animal data were analyzed by analysis of variance (ANOVA), with P≤0.05 considered significant (Prism software; GraphPad, La Jolla, CA) and variables with significant effects by treatment group were subjected to pairwise mean comparisons using the Tukey-Kramer test. Data associated with this study are available for download from the USDA Ag Data Commons at XXX and the phylogenetic analyses from https://github.com/flu-crew/datasets accessed on Date XXX.

## ACNOWLEDGEMENTS

We thank producers, swine veterinarians, and veterinary diagnostic laboratories for participating in the USDA Influenza A Virus in Swine Surveillance System. The authors thank Michelle Harland, Nicholas Otis, and Katharine Young for technical assistance with laboratory techniques, Jason Huegel, Justin Miller, Randy Leon, Alyssa Dannen, and Adam Hartfiel for animal care. This study was supported by USDA-ARS, USDA-APHIS, and the National Institute of Allergy and Infectious Diseases, National Institutes of Health, Department of Health and Human Services contracts HHSN272201400008C and 75N93021C00015. This research used resources provided by the SCINet project of the USDA Agricultural Research Service, ARS project number 0500-00093-001-00-D. JDP was supported in part by an appointment to the USDA-ARS Research Participation Program administered by the Oak Ridge Institute for Science and Education (ORISE) through an interagency agreement between the U.S. Department of Energy and USDA under contract number DE-AC05-06OR23100. Mention of trade names or commercial products in this article is solely for the purpose of providing specific information and does not imply recommendation or endorsement by the U.S. Department of Agriculture, DOE, or ORISE. USDA is an equal opportunity provider and employer.

## SUPPLEMENTAL FIGURES

**Supplemental Figure 1.**
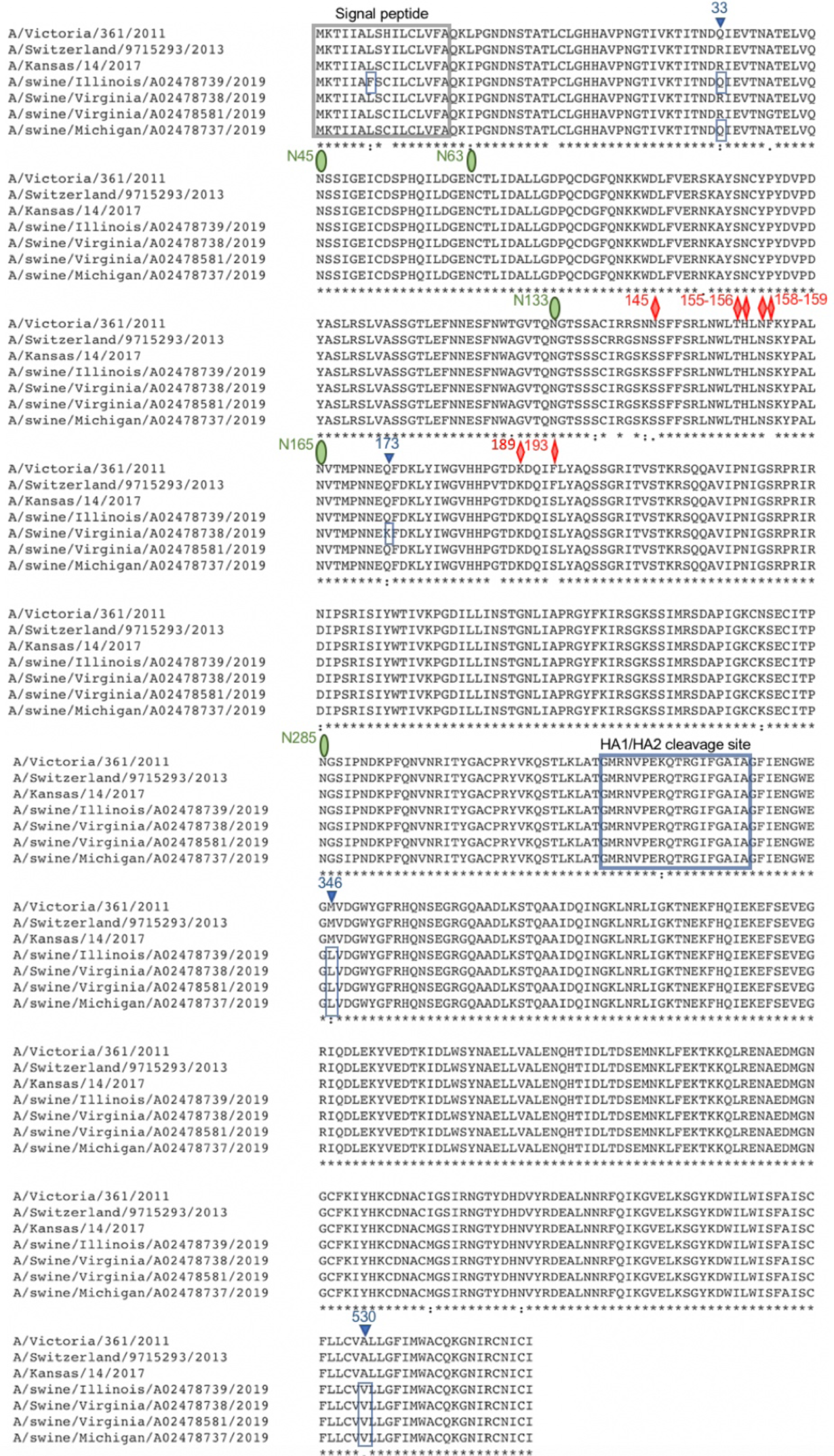
Hemagglutinin alignment of four swine spillover detections in Virginia, Illinois, and Michigan compared to that of three human seasonal influenza vaccine strains representing evolution of the 3c3.A clade from 2011 to 2017. Amino acid differences between spillover strains to that of A/Kansas/14/2017 (blue triangles) are noted within the signal peptide and at positions 33, 173, 346, 530 according to the numbering scheme of Mastrovich et al. (47). The positions of predicted HA1 glycosylation sites (green oval) and the seven key human H3 antigenic residues (red diamonds) previously defined (48, 49) are noted.

**Supplemental Figure 2.**
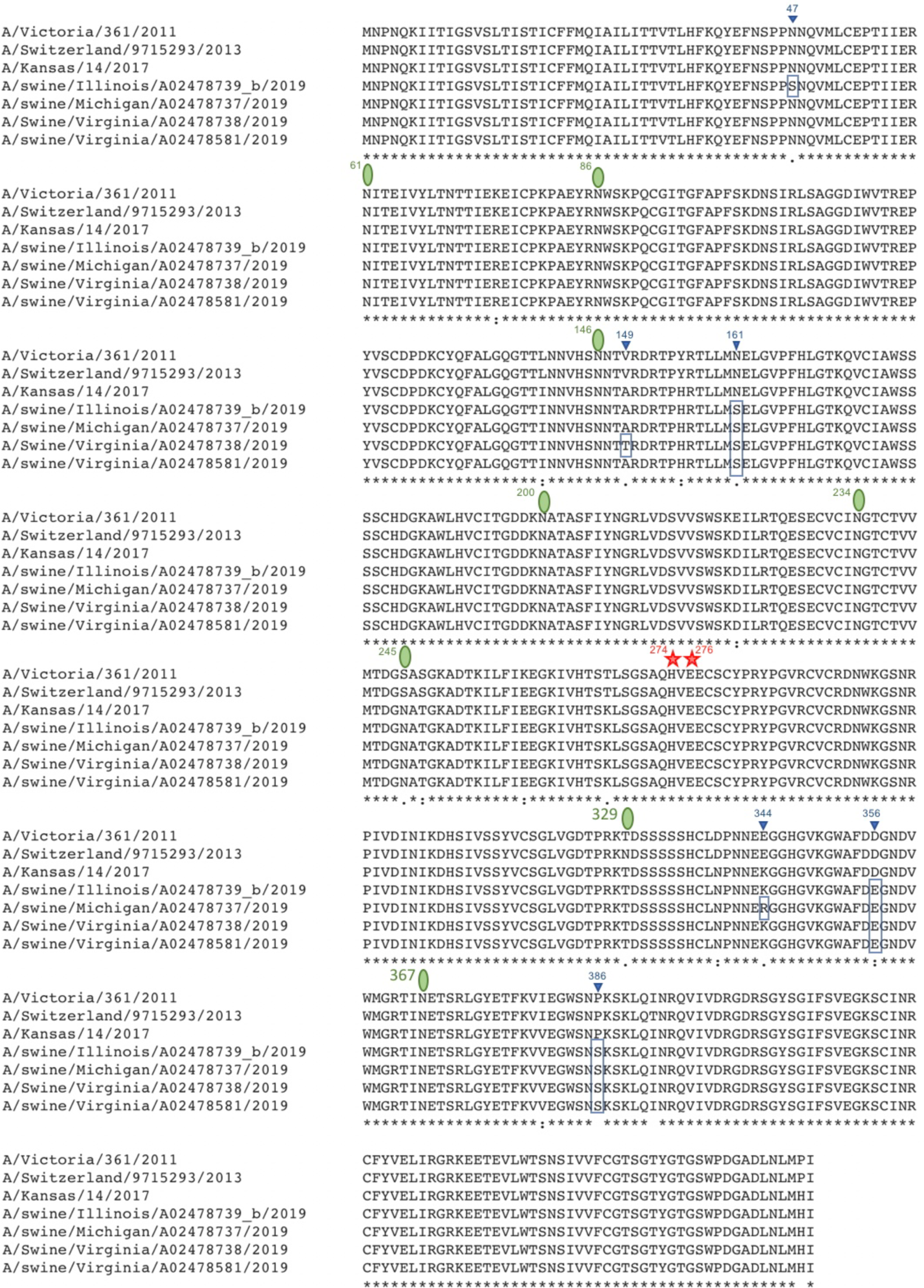
Neuraminidase alignment of four swine spillover detections in Virginia, Illinois and Michigan compared to that of three human seasonal influenza vaccine strains representing evolution of the 3c3.A clade from 2011 to 2017. Amino acid differences between spillover strains to that of A/Kansas/14/2017 (blue triangles) are shown. The positions of key N2 glycosylation sites (green oval) (50) and two key amino acids within the sialic acid binding domain associated with Oseltamivir resistance (red star) are noted.

